# Cross-reactive macaque antibodies targeting marburgvirus glycoprotein induced by multivalent immunization

**DOI:** 10.1101/2023.09.23.559102

**Authors:** Benjamin M. Janus, Ruixue Wang, Thomas E. Cleveland, Matthew C. Metcalf, Aaron C. Lemmer, Kenneth Class, Thomas R. Fuerst, Gilad Ofek

**Affiliations:** Department of Cell Biology and Molecular Genetics, University of Maryland, College Park, MD, USA; Institute for Bioscience and Biotechnology Research, University of Maryland, Rockville, MD, USA; Biomolecular Measurement Division, National Institute of Standards and Technology, Gaithersburg, MD, USA

## Abstract

We utilized B cells from a Rhesus macaque immunized with a multivalent prime-boost regimen of filovirus antigens to isolate a novel panel of marburgvirus glycoprotein (GP)-specific monoclonal antibodies (mAbs). A heterologous marburgvirus GP probe was used to sort for B cells with cross-marburgvirus reactive breadth. 33 mAbs belonging to 28 unique lineages were expressed and experimentally characterized. Antibody specificities were assessed by binding competition and overlapping pepscan analyses, and were found to map to a previously characterized protective region on GP2 and a cross-filovirus reactive region on GP1, among others. A third of the lineages targeted the predicted receptor binding region (RBR), including two lineages with potent Marburg pseudovirus neutralization that were structurally analyzed and confirmed to recognize this region. Our study describes the discovery and characterization of a diverse panel of antibodies against marburgvirus GP induced by multivalent immunization and provides candidate immunotherapeutics for further study and development.

**Author Summary:** Marburgviruses were the first filoviruses characterized to emerge in humans in 1967, and have led to multiple outbreaks since then with average case fatality rates of ∼50%. Although a vaccine and monoclonal antibody countermeasures have been approved for clinical use against the related Ebola viruses, these are ineffective against marburgviruses or other filoviruses. As such, gaps exist in the clinical toolkit against filoviruses, in particular marburgviruses. Here, we isolated and characterized a novel panel of monoclonal antibodies directed against the marburgvirus surface glycoprotein from an immunized Rhesus macaque. We utilized an antibody isolation method that ensured broad antibody recognition across multiple marburgvirus isolates. Functional and structural analyses revealed that roughly half of the antibodies in the panel mapped to regions on the glycoprotein shown previously to protect from infection, including the receptor binding domain and a protective region on the membrane-anchoring subunit, while a quarter of the antibodies did not fall into any known binding competition group indicating potential novel specificities. Our study advances the understanding of marburgvirus glycoprotein antigenicity and furthers efforts to develop candidate antibody countermeasures against these lethal viruses.

## Introduction

The 2013-2016 Ebola virus (EBOV) outbreak in West Africa, which led to over 28,000 cases and 11,000 deaths, expedited efforts at isolation and development of monoclonal antibody countermeasures against filoviruses [1–16]. Multiple panels of effective antibodies against ebolaviruses have since been isolated, including from human survivors and animal immunizations. Such antibodies invariably target the ebolavirus surface glycoprotein (GP), which mediates virus attachment and entry into host cells. While antibody-mediated virus neutralization represents one of the strongest correlates of protective efficacy, in some cases non-neutralizing antibodies are also protective, through facilitation of immune effector functions [12]. Various domains on ebolavirus GP have been shown to be targeted by protective antibodies, including its base, the GP1-GP2 subunit interface, the internal fusion loop (IFL), the receptor binding region (RBR), the glycan cap, and the membrane proximal external region [11, 12, 17-21]. A clinical trial conducted during the 2018 outbreak of EBOV in the Democratic Republic of the Congo revealed efficacy of two monoclonal antibody therapies, one comprised of a single antibody isolated from a human survivor and the other comprised of a cocktail of three antibodies isolated from immunization of transgenic mice with humanized immunoglobulin genes [6, 13]. Both therapies, which are monospecific to EBOV, have since been approved for clinical use by the Food and Drug Administration [22]. Cross-reactive antibodies that protect against infection by multiple ebolavirus species have also been isolated and are of special interest because they target conserved regions on GP that are thought to be less permissive to viral escape due to functional constraints [8, 9, 11, 18, 23-25]. Such antibodies may also be effective against as yet to be defined pre-emergent species.

Protective antibodies targeting marburgviruses have also been isolated from human survivors and animal immunizations, although in more limited scope [14-16, 26-32]. Compared to antigenic targets on ebolavirus GP, protective mAbs against marburgvirus GP appear to map to fewer antigenic sites, which to date only include the RBR, the N-terminus of GP2 (sometimes referred to as the “GP2 wing”), epitopes within the mucin-like domain, and a target at the base of GP [14-16, 26-32]. Antibodies that target the RBR are generally neutralizing but have thus far only been isolated from survivors of natural infection (or from immunogenetic sequence homologues thereof), suggesting gaps exist in current immunization strategies or antibody isolation strategies for isolation of marburgvirus RBR-specific neutralizing antibodies [14, 31]. Protective antibodies that target the GP2 N-terminus or “wing” have been isolated from natural infection and animal immunizations, but are weakly- or non-neutralizing and are thought to protect by mediating immune effector functions [15, 16]. Antibodies that target the GP mucin-like domain appear to inhibit virus infection by preventing virion budding [30]. While nonhuman primate virus challenge protection studies using various mAbs have shown some success, in particular using RBR-directed neutralizing antibodies isolated from a human survivor, high antibody doses and administration by 5 days post infection appear to be required for protection [33].

With the goal of inducing novel antibody responses against conserved regions on filovirus GPs, we previously explored immune responses in Rhesus macaques immunized with escalating species multivalent prime-boost regimens using antigens of three filovirus species, MARV, Sudan (SUDV), and EBOV [34]. One of the animals, NHP1, developed high titers of neutralizing antibodies against MARV and was chosen for further study [34]. Here, we describe a novel panel of monoclonal antibodies directed against marburgvirus GP isolated from memory B cells of NHP1. Antigen specific single B cells were sorted using a heterologous Ravn (RAVV) marburgvirus GP probe to select for B cells with cross-marburgvirus reactivity. 33 monoclonal antibodies belonging to 28 distinct lineages were characterized and are described herein. Two lineages that belonged to the RBR antigenic group exhibited potent MARV pseudovirus neutralization, representing to our knowledge the first reported isolation of such marburgvirus neutralizing antibodies from animal immunizations.

## Results

### Heterologous probe for isolation of cross-marburgvirus reactive B cells

We previously immunized Rhesus macaques with multivalent combinations of antigens corresponding to three filovirus species that have caused outbreaks in humans, MARV, SUDV, and EBOV [34]. We utilized an escalating species immunization approach that was weighted with MARV and SUDV antigens to augment responses against those species. In one animal (NHP1) that was primed with MARV (*Musoke*) virus-like particles (VLPs) composed of GP, VP40, and NP, and then boosted with a bivalent mixture of MARV (*Musoke*) and SUDV (*Yambio*) VLPs, followed by two trivalent boosts with recombinant MARV (*Angola*), SUDV (*Yambio*), and EBOV (*Mayinga*) GPΔMuc proteins, high titers of marburgvirus neutralizing antibodies were detected (**Fig. 1A,C**) [34].

**Figure 1.**
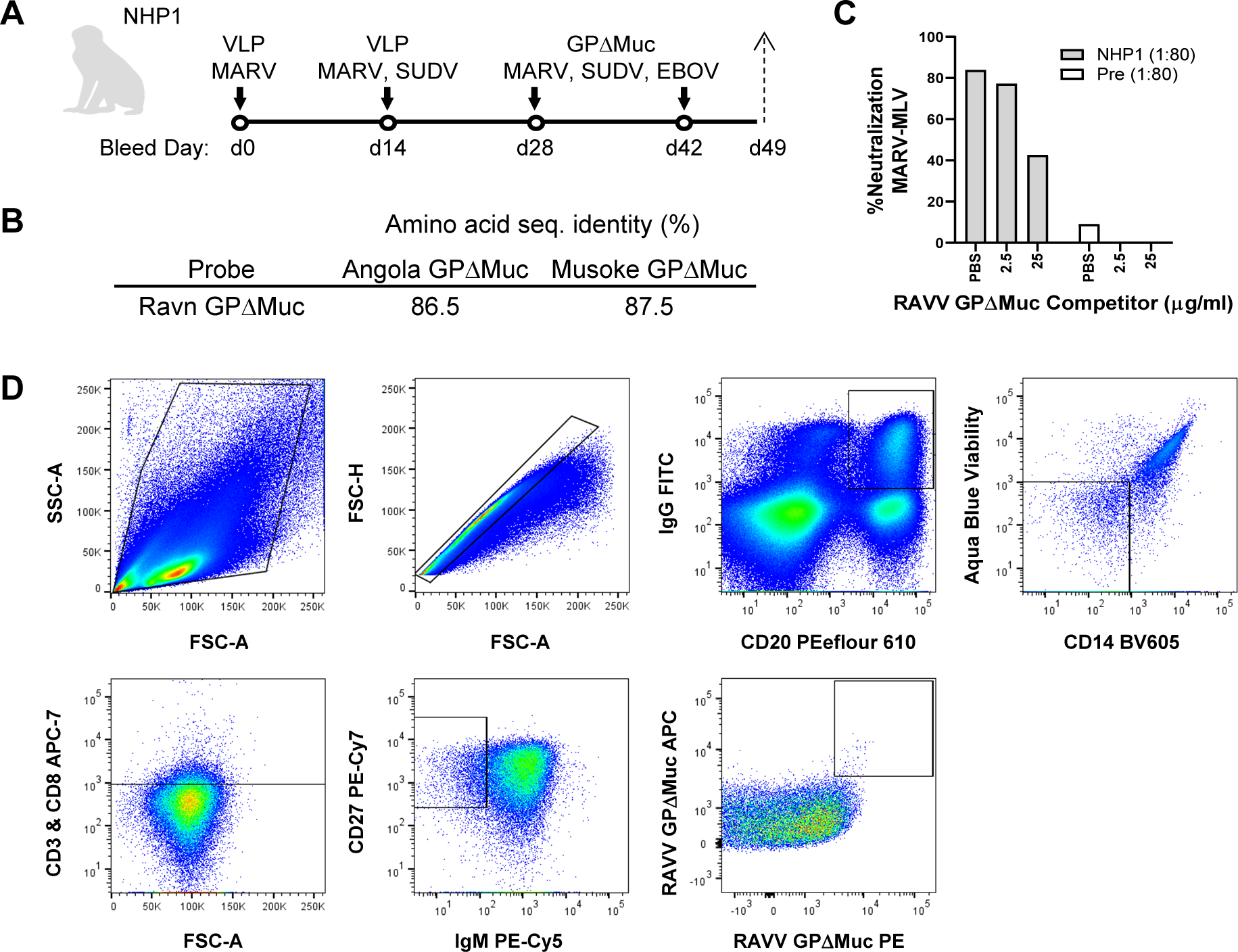
Heterologous marburgvirus GP-specific antibodies isolated from an immunized Rhesus macaque. **A.** Species escalating multivalent prime-boost immunization regimen administered to NHP1 whose day 49 PBMCs were used for antigen specific single B cell sorting. MARV VLPs were based on strain *Musoke* and recombinant MARV GPΔMuc boosts were based on strain *Angola*. **B.** Sequence divergence of the heterologous RAVV GPΔMuc B cell sorting probe from autologous *Musoke* and *Angola* MARV immunogens. **C.** Validation of heterologous RAVV GPΔMuc probe as a competitor for NHP1 serum neutralization of MLV-MARV. **D.** Sorting of RAVV GPΔMuc double positive memory B-Cells with the phenotype CD27+, IgG+, CD20+, Aqua Dead-, CD3-, CD8-, CD14-, IgM-.

With the goal of isolating cross-marburgvirus reactive antibodies we sought to develop a probe that could be used to select for cross-marburgvirus reactive memory B cells from animal NHP1. Towards this end, we utilized a recombinant mucin-deleted ectodomain based on the sequence of Ravn virus, RAVV GPΔMuc, that was previously shown to be recognized by NHP1 serum and that diverged in amino acid sequence from autologous *Musoke* and *Angola* MARV GPs by ∼12-15% (**Fig. 1B**) [34]. To further validate RAVV GPΔMuc as a probe for B cell isolation, we used it as a competitor in MARV-MLV pseudoparticle neutralization assays to assess the presence of RAVV GPΔMuc-reactive heterologous neutralizing antibodies in NHP1 serum. As shown in **Fig. 1C**, pre-incubation of 25 μg/ml of RAVV GPΔMuc probe with NHP1 serum prior to addition to pseudoviruses and target cells successfully reduced serum neutralization by ∼60%. These results confirmed the presence of heterologous cross-marburgvirus reactive neutralizing antibodies in NHP1 serum and validated the use of RAVV GPΔMuc as a heterologous probe for B cell sorting.

### Antigen specific memory B cell sorting

To isolate RAVV GPΔMuc reactive B cells, we stained NHP1 terminal bleed (day 49) peripheral blood mononuclear cells (PBMCs) with avi-tag biotinylated RAVV GPΔMuc protein conjugated with two versions of fluorescently labeled streptavidin, APC and PE, along with a cocktail of reagents targeting memory B cell surface markers to ensure selection of B cells that were of the phenotype IgG^+^IgM^-^CD20^+^CD14^-^CD3^-^CD8^-^CD27^+^, RAVV GP^++^ (**Fig. 1D**) [35, 36]. Of ∼480 wells of cells that were collected, we utilized the first 96 wells to establish an initial panel of monoclonal antibodies through nested PCR amplification of heavy and light chain antibody variable regions [35, 36]. 58 out of 96 wells yielded successful amplification of both heavy and light chain antibody products and 34 pairs were selected for experimental characterization. Heavy and light chains variable regions were synthesized and subcloned into mammalian human IgG1 expression vectors for transient expression in HEK293F cells. 33 out of the 34 antibodies expressed to sufficient levels to permit experimental characterization.

Sequence analysis revealed that the 33 antibodies belonged to diverse immunogenetic backgrounds, corresponding to roughly 10 IGHV and 21 IGLV genes (**Fig. 2A**). A majority of the heavy chains were of VH3-background (**Fig. 2A**). Further analysis of the sequences revealed rates of somatic mutation that ranged from 0.7-12.3% and 1.0-9.7% for heavy and light chains respectively. Heavy chain CDR3 loop lengths ranged from 6 to 20 amino acids (**Fig. 2B**). A majority of the antibodies in the panel represented independent clonotypes, although 9 variants belonged to one of four shared lineages (**Fig. 3A**).

**Figure 2.**
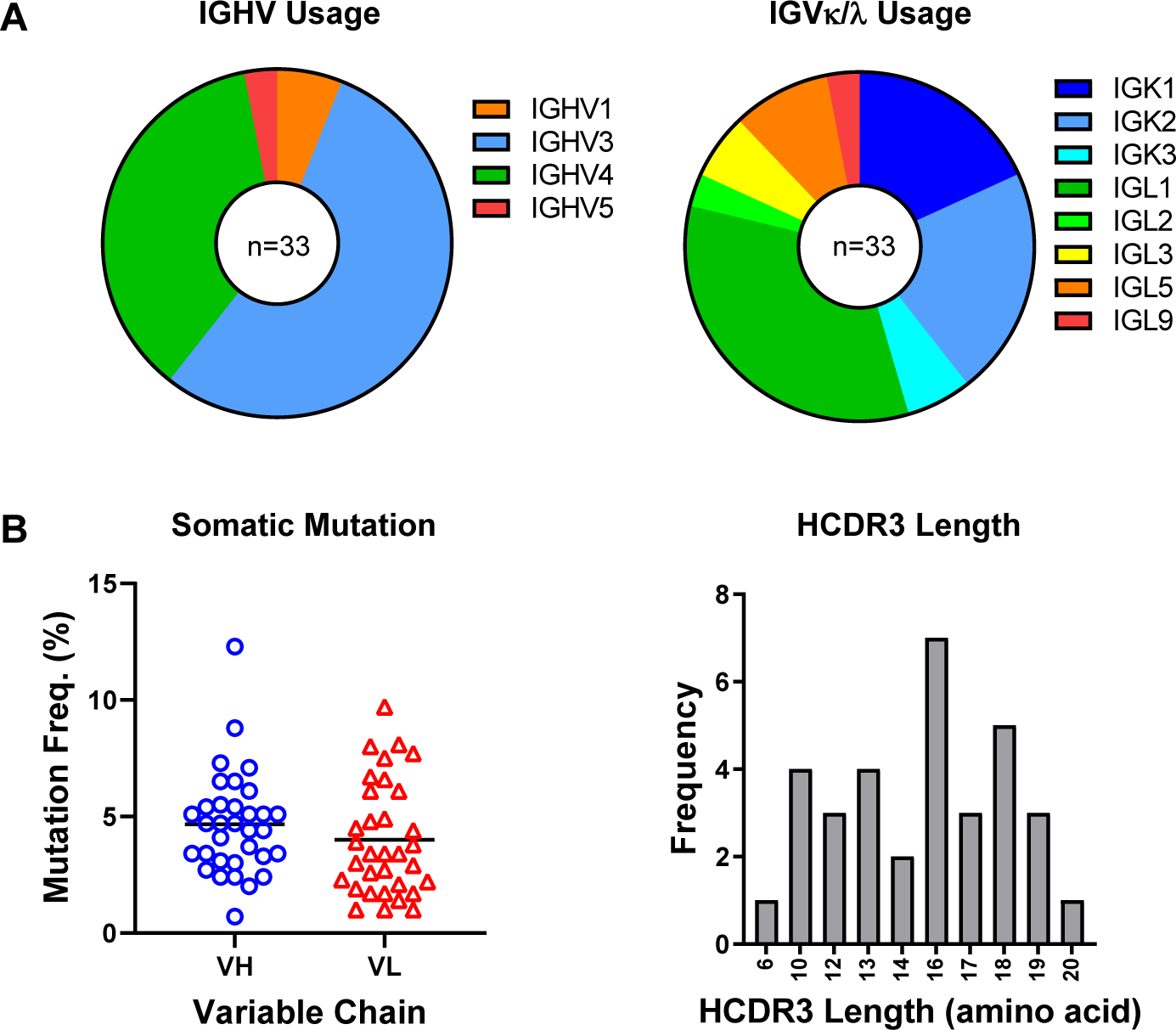
Antibody panel heavy and light chain sequence features. **A.** Heavy and light chain gene usage of the 33 expressed monoclonal antibodies. **B.** Heavy chain and light chain somatic mutation frequencies, shown as percentages of total amino acids in variable regions. **C.** Heavy chain CDR3 lengths

**Figure 3.**
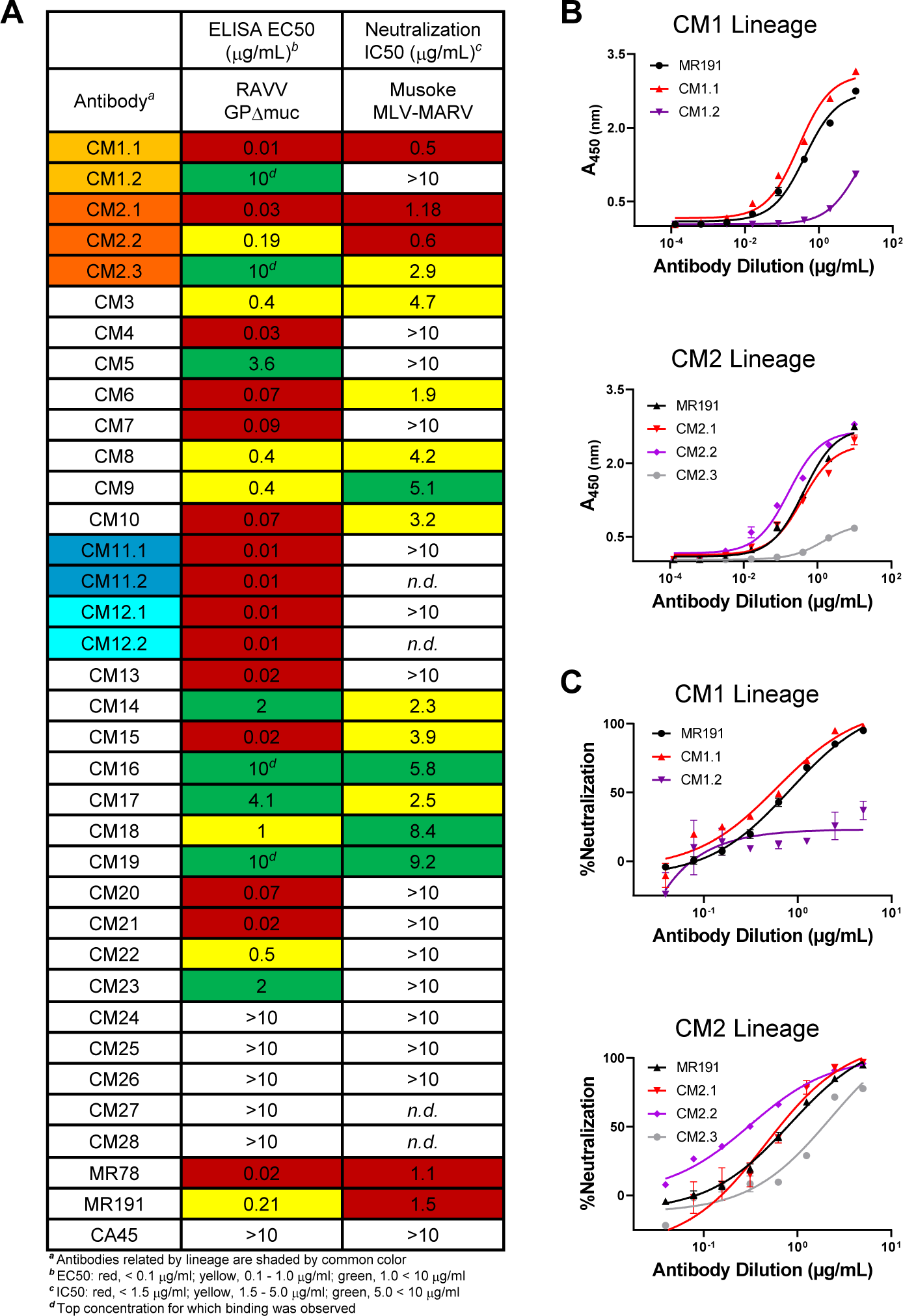
GP binding and pseudovirus neutralization. **A.** Antibody ELISA binding EC50s to RAVV GPΔMuc and neutralization IC50s against MARV-GP MLV pseudotyped particles. **B.** RAVV GPΔMuc ELISA binding of CM1 and CM2 lineage variants. **C.** Neutralization of MARV-GP pseudoviruses by CM1 and CM2

### GP binding and psuedoparticle neutralization

We next assessed the binding of the 33 antibodies to RAVV GPΔMuc by ELISA, alongside control marburgvirus GP-specific antibodies MR78 and MR191 and ebolavirus GP-specific antibody CA45 [14, 18]. As shown in **Fig. 3A**, 28 of the novel antibodies (roughly 85%) bound RAVV GPΔMuc with EC50 values that ranged from 0.01-10 μg/ml, confirming that our B cell sorts led to successful isolation of marburgvirus GP-specific monoclonal antibodies. Indeed, some of the antibodies bound with EC50 values that were commensurate or better than those observed for antibodies MR78 and MR191 (**Fig. 3A**).

Using a maximum antibody concentration of 10 μg/ml, we tested the 28 GP-reactive antibodies for the capacity to neutralize Murine Leukemia Virus (MLV) particles pseudotyped with MARV GP (*Musoke*) [37]. 16 of the 28 antibodies tested (∼57%) exhibited neutralization of MLV-MARV pseudoviruses to different degrees, with neutralization IC50 values that ranged from 0.5-9.2 μg/ml (**Figs. 3A**). Two of the antibody lineages, CM1 and CM2, exhibited the most potent neutralization observed in the panel with IC50s that ranged from 0.5-1.8 μg/ml, on par with IC50’s obtained for the MR191 and MR78 controls (**Figs. 3A,C**) [14]. While some of the variants that belonged to the CM1 and CM2 lineages exhibited weak or undetectable neutralization, namely mAbs CM1.2 and CM2.3, such differences correlated with differences in binding capacity to recombinant GP by ELISA (**Fig. 3B**). We also note that mAbs CM1.2 and CM2.3 expressed at low levels and were prone to proteolytic cleavage, consistent with potential biochemical instability. Nonetheless, our results confirm that a majority of the isolated mAbs were able to effectively recognize heterologous RAVV GPΔMuc, with two novel lineages exhibiting potent MARV pseudovirus neutralization.

### Cross-filovirus GP recognition

Although the main goal of the present study was to isolate cross-marburgvirus reactive antibodies, since animal NHP1 received multivalent immunizations that included SUDV and EBOV based antigens, we next assessed whether any of the 28 antibody lineages had the capacity to recognize ebolavirus GPs as well. Towards this end, we first tested the mAbs in the panel for recognition of recombinant EBOV GPΔMuc by ELISA. While a majority of the antibodies had weak or undetectable binding to EBOV GPΔMuc (not shown), two antibodies – CM16 and CM20 – did exhibit detectable EBOV GPΔMuc binding (**Fig. 4**). Testing of CM16 and CM20 for recognition of other ebolavirus GPs, namely SUDV and BDBV GPΔMuc, revealed that both antibodies recognized SUDV and BDBV GPΔMuc equally well if not better than their recognition of EBOV GPΔMuc (**Fig. 4**). Binding of CM16 and CM20 to ebolavirus GPs was similar to that observed for the control ebolavirus-specific antibody CA45, although their recognition of RAVV GPΔMuc trailed that of the MR191 control (**Fig. 4**).

**Figure 4.**
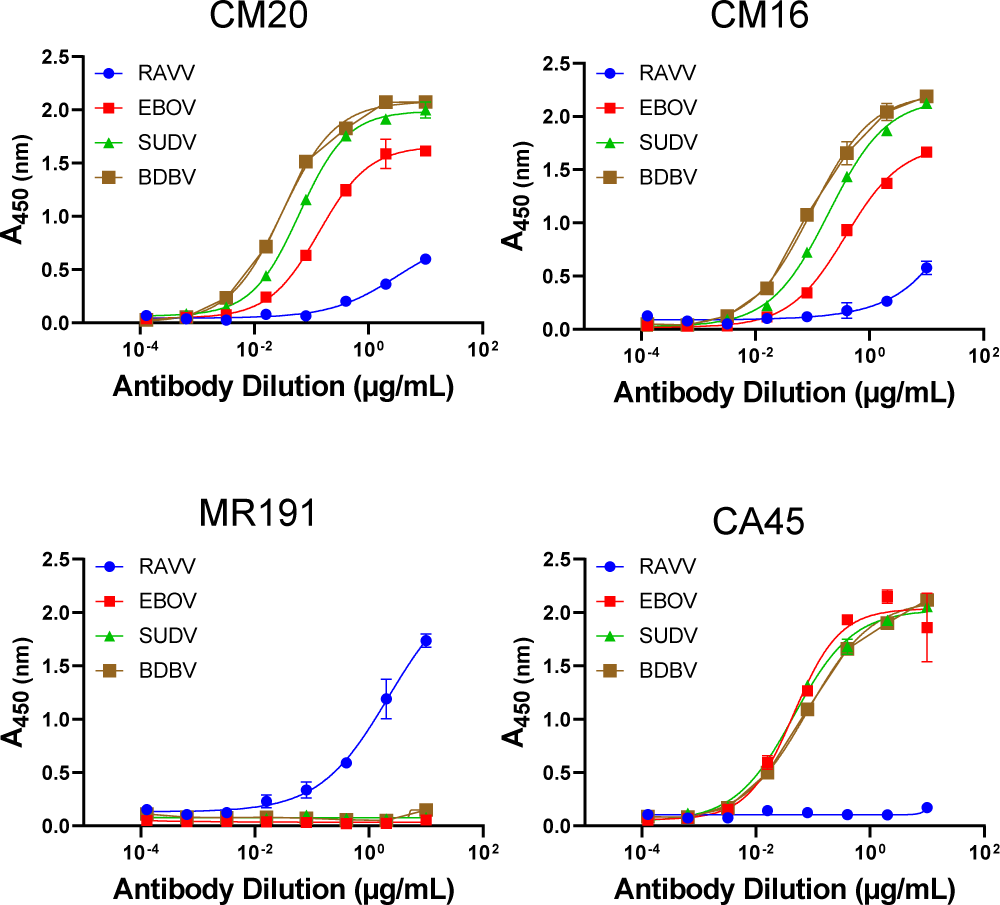
Cross-filovirus GP recognition. ELISA binding profiles of antibodies CM16 and CM20 to RAVV, EBOV, SUDV, and BDBV GPΔMuc proteins. Marburgvirus-specific mAb MR191 and ebolavirus-specific cross-reactive mAb CA45 are shown as controls.

While mAbs CM16 and CM20 exhibited only weak or undetectable neutralization of MARV GP pseudoviruses, in view of their observed cross-filovirus GP recognition we assessed their neutralization capacity against ebolaviruses. Neither antibody exhibited strong neutralization of EBOV, SUDV, or BDBV pseudoviruses, suggesting that they either target a conserved epitope that does not confer inhibition of entry or that other features rendered them ineffective in preventing viral entry at the concentrations used in these assays. Taken together, our results nonetheless confirm that the escalating species multivalent prime-boost immunization regimen given to NHP1 led to successful induction of monoclonal antibodies with cross-filovirus reactive breadth.

### Epitope mapping by overlapping pepscan analysis

Prior to undertaking overlapping pepscan analysis for epitope mapping, we first assessed whether any of the GP-reactive antibodies in the panel could recognize a contiguous, non-conformational epitope on RAVV GPΔMuc. Towards this end, RAVV GPΔMuc protein was applied to a denaturing SDS-PAGE gel and subjected to standard Western blotting procedures, using each individual GP-reactive antibody as a probe. Five of the tested mAbs gave detectable signal by Western blot analysis (**Fig. 5A**). Two mAbs, CM13 and CM21, reacted with a band corresponding to the size of GP1, while the remaining three mAbs, CM10, CM11.1, and CM12.1, targeted a band corresponding to the size of GP2 (**Figs. 5A**).

**Figure 5.**
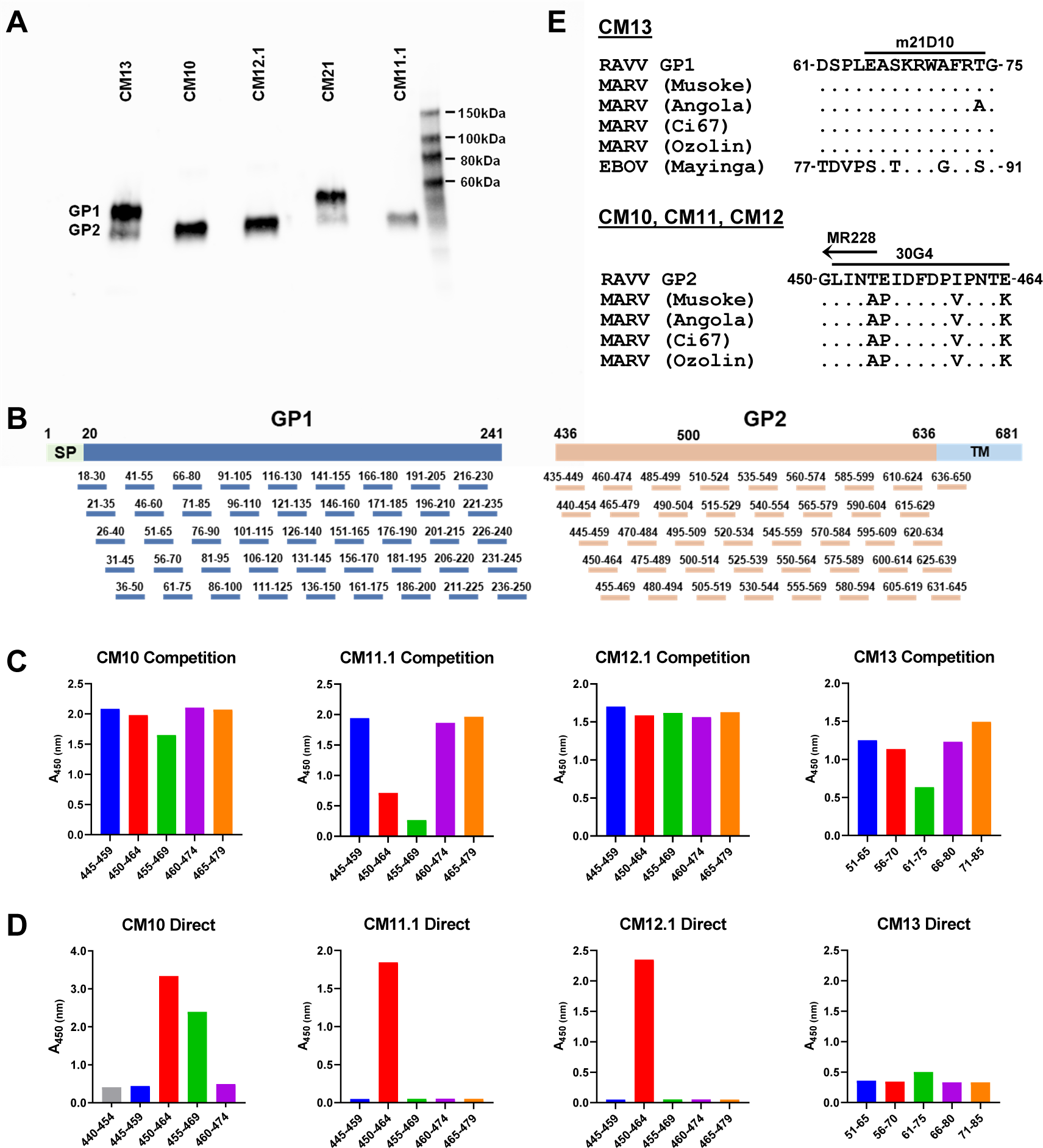
Epitope mapping by overlapping pepscan analysis. **A.** SDS-PAGE Western blots of RAVV GPΔMuc probed with five mAbs observed to give detectable recognition of denatured GP. **B.** Schematic of overlapping 15-mer peptides across the GP1 and GP2 subunits of RAVV GPΔMuc used for pepscan analyses. **C.** Overlapping peptide ELISA binding competition for RAVV GPΔMuc recognition by mAbs CM10, CM11.1, CM12.1, and CM13, focused on regions of GP1 and GP2 that exhibited competition. **D.** Direct binding of mAbs CM10, CM11.1, CM12.1, and CM13 to overlapping peptides, focused on regions defined in *C*. **E.** Sequence alignments across marburgviruses of the GP1 epitope of mAb CM13, top, and the GP2 epitope of mAbs CM10, CM11, and CM12, bottom. Overlapping epitopes of previously characterized mAbs m21D10, 30G4, and MR228 are shown as bars above.

To further map the epitopes of these five Western-blot reactive mAbs, we generated a panel of overlapping 15-mer peptides covering the sequences of both GP1 and GP2 of RAVV GPΔMuc, and undertook both competition and direct ELISA binding analyses (**Fig. 5B-D**). For mAbs CM10, CM11.1, and CM12.1, which were predicted to target the GP2 subunit by Western blot analysis, our analysis focused on binding to 46 overlapping peptides covering the GP2 ectodomain, namely spanning residues 435 to 650. To assess whether any of the 46 overlapping GP2 peptides could successfully compete for mAb recognition of RAVV GPΔMuc we co-incubated each mAb with each individual peptide and then added the mixture to ELISA wells coated with RAVV GPΔMuc. These assays revealed that peptides within the GP2 N-terminus (or “wing”) corresponding to residues 450-464 and 455-469, a region previously shown to be the target of protective antibodies against marburgviruses, successfully competed for CM10 and CM11.1 recognition of RAVV GPΔMuc (**Fig. 5C,E**) [15, 16]. None of the 46 overlapping GP2 peptides effectively competed for CM12.1 mAb recognition of RAVV GPΔMuc (**Fig. 5C**). To further confirm CM10 and CM11.1 recognition of the GP2 N-terminus, and map the epitope of mAb CM12.1, we undertook direct ELISA binding analyses using the same overlapping peptides spanning GP2 residues 440-479. mAbs CM10 and CM11.1 both bound directly to peptide 450-464, while mAb CM10 also exhibited biding to peptide 455-469. Notably, mAb CM12.1 also bound to peptide 450-464 directly, despite the inability of this peptide to effectively compete with RAVV GPΔMuc for CM12.1 binding (**Fig. 5C,D**). Thus, mAbs CM10, CM11.1, and CM12.1 target an overlapping epitope within the GP2 N terminus, one that overlaps with epitopes of previously reported protective marburgvirus mAbs isolated from natural infection and animal immunizations (**Fig. 5E**) [14-16, 38].

Next, to map the epitopes of GP1 Western-blot reactive mAbs CM13 and CM21 we employed a similar strategy, but utilized a set of overlapping GP1 peptides instead (**Fig. 5B**). 45 overlapping 15-mer peptides spanning GP1 ectodomain residues 18-250 were used as competitors for CM13 and CM21 binding to RAVV GPΔMuc (**Fig. 5B,C**). Binding of CM13 to RAVV GPΔMuc was competed ∼50% by a peptide spanning GP1 residues 61-75, although direct recognition of this peptide was weaker (**Fig. 5C,D**). We note that peptide 61-75 lies in the vicinity of the predicted RBR on GP1, and partially overlaps with the epitope of a previously reported pan-filovirus reactive murine antibody, m21D10, that was also isolated from multivalent immunization [39]. In contrast to CM13, none of the 45 overlapping 15-mer GP1 peptides competed with mAb CM21 for binding to RAVV GPΔMuc, nor were any recognized by direct ELISA (not shown), indicating other means will be necessary to map its epitope.

### Determination of antigenic binding competition groups

To further classify the antigenic targets of the antibodies in the panel, we undertook GP binding competition analyses to define antigenic competition groups. Four antibodies were selected as antigenic benchmarks for recognition of RAVV GPΔMuc against which all antibodies in the panel were tested as competitors. Benchmark mAbs included CM10 and CM13 that were mapped to continuous epitopes on GP2 and GP1, respectively, along with two potent MARV neutralizing antibodies, CM1.1 from the present study and antibody MR191, a previously reported neutralizing antibody that targets the predicted RBR on GP (**Fig. 6**) [14]. Our binding competition assay entailed pre-incubation of each GP-reactive antibody in the panel with biotinylated RAVV GPΔMuc for 1 hour followed by addition of the complex to ELISA plates pre-coated with each of the four antigenic benchmark antibodies. The degree to which the benchmark antibodies could capture biotinylated RAVV GPΔMuc alone or in the presence of competitor antibodies in the panel was assessed by detection with HRP-conjugated anti-biotin antibody.

**Figure 6.**
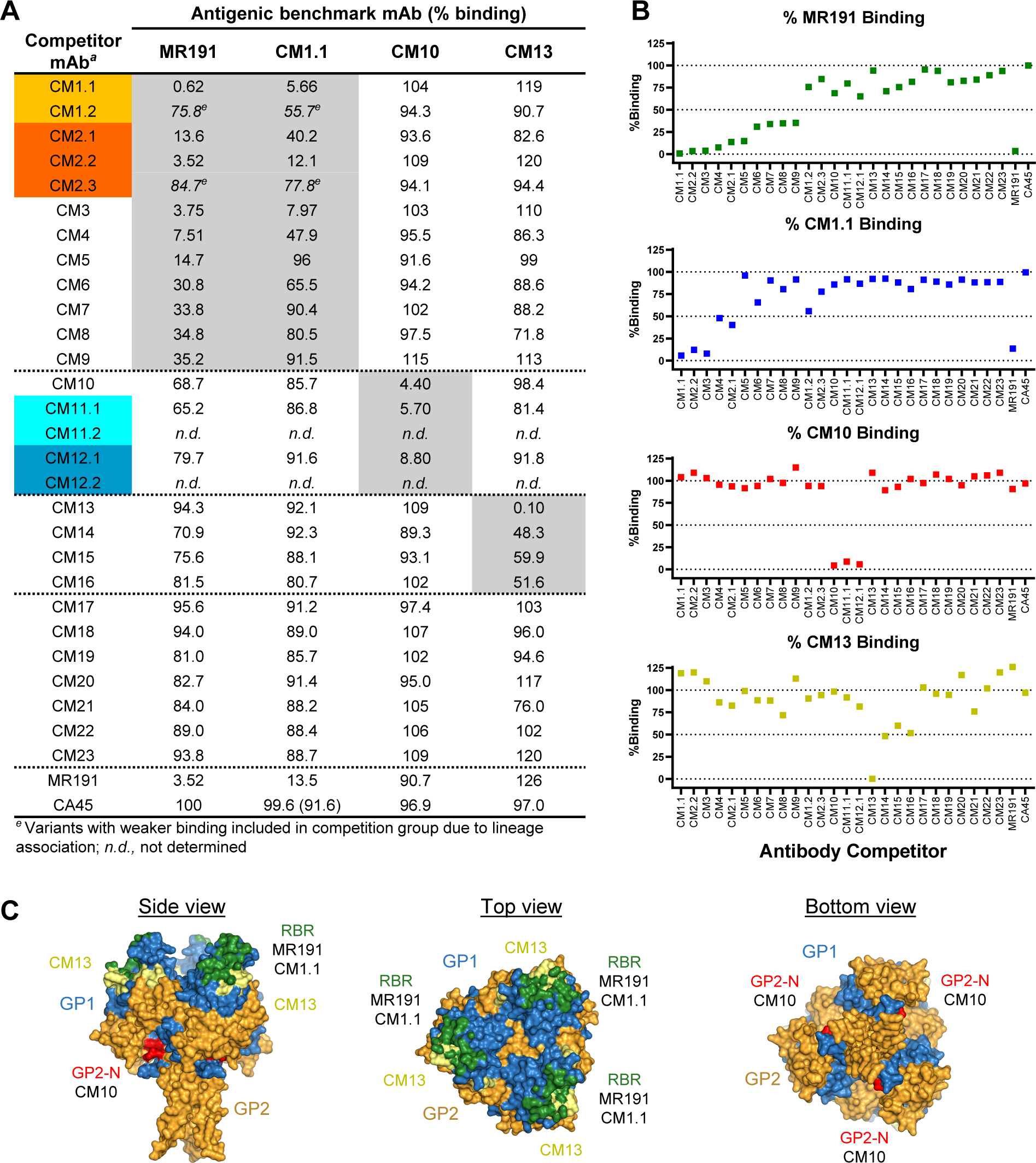
Antibody binding competition groups. **A.** Shown are %-binding values of each antigenic benchmark mAb to biotinylated RAVV GPΔMuc in presence of each competitor mAb. Percentages are calculated relative to capture of biotinylated RAVV GPΔMuc in the absence of a competing mAb. Ebolavirus-specific mAb CA45 was used as a negative control. **B.** Graphical representation of the %-binding competition results shown in *A*. **C.** Epitopes of antigenic benchmark mAbs MR191, CM1.1, CM10, and CM13, mapped onto a surface of RAVV GPΔMuc (PDB ID 6BP2), shown in three orientations with GP1 and GP2 colored blue and orange, respectively. The predicted RBR is colored green, based on the epitope of MR191. The first ordered residues within the N terminus of GP2 (GP2-N) are colored red and the CM13 epitope is colored yellow.

As shown in **Fig. 6A,B**, roughly 30% of the antibodies in the panel (10 out of 33), fell within the RBR antigenic competition group in that they blocked between 64% to greater than 99% of MR191 binding to RAVV GPΔMuc. Two potent neutralizing antibodies, CM1.1 and CM2.1, and a subset of their lineage mates, fell within this MR191-directed competition group as well (**Fig. 6A,B**). Indeed, antibody CM1.1 which was also used as an antigenic benchmark mAb itself, was the most effective MR191 competitor of all the antibodies tested, knocking out more than 99% of MR191 binding when pre-incubated with RAVV GPΔMuc, better than MR191’s competition against itself (**Fig. 6A,B**). Remarkably, out of the ten antibodies that effectively competed away more than 65% of MR191’s binding to GP, only four effectively competed with benchmark antibody CM1.1, namely, CM2.2, CM3, CM4, and CM2.1 (**Fig. 6A,B**). The remaining five MR191 competitors, CM5, CM6, CM7, CM8, and CM9, competed to a lesser degree or not at all with CM1.1, suggesting that CM1.1 binding to GP was more difficult to block than MR191 or that the MR191 epitope coincided more directly with these five antibodies. While antibodies CM1.2 and CM2.3 were not effective at competing with either MR191 or CM1.1, despite belonging to the CM1 and CM2 lineages, we note that these two variants exhibited substantially lower expression yields, were prone to proteolytic cleavage, and did not bind RAVV GPΔMuc as effectively as other lineage members suggesting they were biochemically unstable (**Figs. 6A,B and 3B**).

For antigenic benchmark antibody CM10 which was shown above to be specific to the GP2 N terminus by overlapping pepscan analysis (**Fig. 5**), the assay revealed as expected that pre-incubation of GP with antibodies CM11.1 and CM12.1 reduced CM10 recognition of GP by 94% and 91%, respectively (**Fig. 6A,B**). CM11.1 and CM12.1, like CM10, bound denatured GP by Western blot and were mapped to the same overlapping peptides within the GP2 N terminus as CM10 (**Fig. 5**). Thus, the present binding competition assays confirmed that all three antibodies, CM10, CM11.1, and CM12.1 fell within the same antigenic competition group and could recognize an overlapping epitope within the N terminus of GP2 in the context of the trimeric GP ectodomain. Two lineage mates of CM12.1 and CM11.1, CM12.2 and CM11.2, respectively, were not tested in these assays, but were confirmed in a parallel study to target the same GP2 epitope (B. Janus, G. Ofek, unpublished results). None of the other antibodies in the panel successfully competed with CM10 for GP recognition (**Fig. 6A,B**).

Lastly, for antigenic benchmark antibody CM13 that targets a contiguous pan-filovirus epitope on GP1 in the vicinity of the RBR, the binding competition assays revealed that three other antibodies fell within its antigenic competition group: CM14, CM15, and CM16. Pre-incubation of RAVV GPΔMuc with any one of the three antibodies blocked CM13 recognition of GP by 40-52% (**Fig. 6A,B**). None of these three antibodies recognized denatured GP by Western-blot analysis, suggesting their epitopes were conformational, or incompatible with the overlapping peptide residue boundaries tested by pepscan analysis. We note that antibody CM21, which we could not map by pepscan analysis but appears to recognize denatured GP1 by Western-blot analysis, blocked CM13 binding to GP by ∼25%, indicating possible overlap in their epitopes (**Figs. 5 and 6A,B**).

Our binding competition assays also revealed that pre-incubation of RAVV GPΔMuc with several antibodies in the panel could enhance benchmark antibody binding to GP. In particular, mAbs CM1.1 and CM2.2 within the RBR-directed antigenic competition group, enhanced the binding of benchmark mAb CM13 to RAVV GPΔMuc by ∼20% (**Fig. 6A,B**). Since antibody cooperativity in virus neutralization has been reported for ebolaviruses, further studies will be necessary to assess if cooperativity in binding observed here also translates into cooperativity in virus neutralization [9, 40].

The seven remaining GP-reactive antibodies in the panel, CM17 through CM23 did not robustly fall into any one of the four antigenic competition groups tested (**Fig. 6A,B**). These antibodies may target epitopes on RAVV GPΔMuc distinct from those of the benchmark antibodies used, although we cannot exclude the possibility that the absence of effective competition is a result of insufficient binding affinity to RAVV GPΔMuc.

### Analysis of CM1.1 and CM2.1 recognition of RAVV GPΔMuc by negative stain electron microscopy

To confirm the GP binding targets of the two antibody lineages that exhibited the highest virus neutralization potency, CM1 and CM2, we analyzed their recognition of RAVV GP by negative stain electron microscopy (NSEM). Towards this end, the fragments of antigen binding (Fabs) of CM1.1 and CM2.1 were expressed and individually complexed with recombinant RAVV GPΔMuc protein. Each complex was applied to EM grids and stained with uranyl formate prior to imaging using a Talos Arctica (200 kV) system. Data processing and 2D classification were performed using RELION [41]. As shown in **Fig. 7A,B**, 2D classes generated for both the CM1.1 Fab:RAVV GPΔMuc and CM2.1:RAVV GPΔMuc complexes yielded particles with either one or two Fabs bound at the apex of GP. Observed structures were consistent with those observed for antibodies that target the predicted marburgvirus GP RBR, specifically antibody MR191 that is shown in **Fig. 7C** for comparison [14, 42].

**Figure 7.**
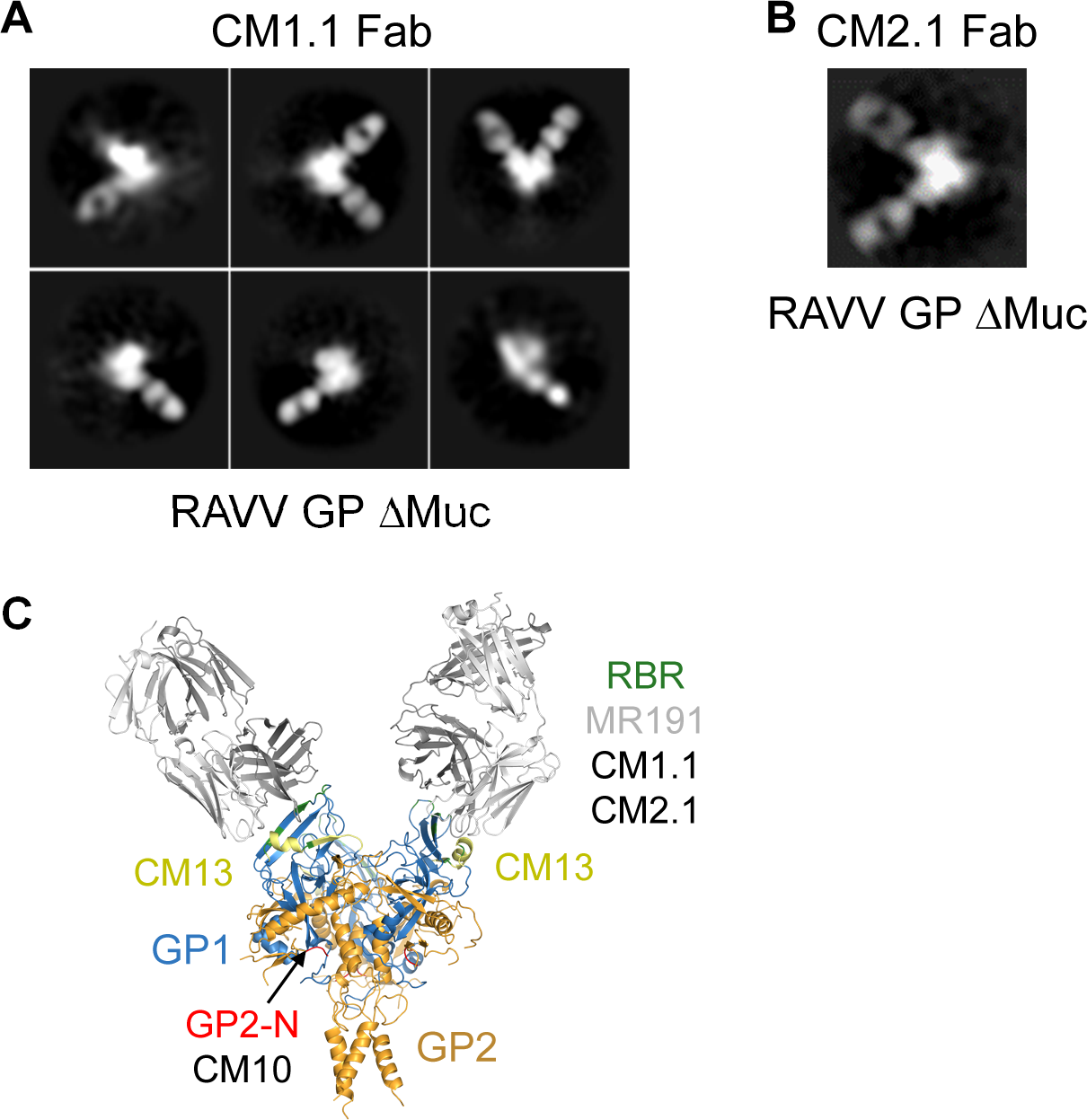
CM1.1 and CM2.1 recognition of GP by negative stain electron microscopy. Negative stain 2D classes of the fragments of antigen binding of antibodies CM1.1 (**A**) and CM2.1 (**B**) in complex with recombinant RAVV GPΔMuc protein. **C.** Crystal structure of RAVV GPΔMuc (PDB 6BP2) colored as in **Fig. 3C**, in complex with RBR-directed neutralizing antibody MR191, gray, shown for comparison.

## Discussion

We isolated and characterized a novel panel of monoclonal antibodies directed against marburgvirus GP from a nonhuman primate previously immunized with an escalating species multivalent regimen of filovirus immunogens. To select for B cells with cross-marburgvirus reactivity, antigen specific single memory B cell sorting was undertaken using a heterologous probe based on the sequence of RAVV GP. 58 pairs of heavy and light chains were successfully amplified, and after downselection 33 mAbs belonging to 28 unique clonotypes were expressed and experimentally characterized. 23 of the 28 clonotypes bound to recombinant RAVV GPΔMuc by ELISA, with 13 of the clonotypes exhibiting varying degrees MARV pseudovirus neutralization.

GP pepscan binding analyses combined with antibody binding competition assays revealed that roughly 70% of the GP-reactive clonotypes (16 of the 23) fell into one of three distinct antigenic groups, which included the RBR, a protective region within the N terminus of GP2, and a contiguous pan-filovirus target on GP1. The remaining 30% of clonotypes could not be mapped by the assays used, suggesting they may possess novel specificities or that the assays were not sufficiently sensitive in those cases.

Within the RBR antigenic group, two antibody lineages, CM1 and CM2, exhibited the highest neutralization potency against MARV obtained in the study. Notably, the neutralization IC50 of mAb CM1.1 was as low as 0.5 μg/ml which was more potent than the IC50s observed for controls MR78 and MR191 in our assays. To our knowledge, other than a panel of antibodies isolated alongside MR78 and MR191 from a human survivor of MARV infection, and associated bioinformatic homologues thereof, these are the only other cases of reported neutralizing antibodies shown to target the marburgvirus RBR, and represent the first such mAbs isolated from immunization [14, 31]. It remains to be determined whether the induction and isolation of the CM1 and CM2 lineages was a result of the escalating species multivalent prime-boost immunization regimen administered to animal NHP1 or alternatively to features of the downstream single B cell sorting pipeline that was utilized to isolate the panel as a whole, including the heterologous RAVV GPΔMuc probe.

Three antibody lineages in the panel, CM10, CM11, and CM12, mapped to the N terminus of GP2, a region previously shown to be a target of non-neutralizing but protective antibodies [15, 16]. CM10, CM11, and CM12 all recognized a continuous epitope within this region, partially or fully overlapping with epitopes of previously characterized protective mAbs [15, 16]. The mapped epitopes of all three lineages were fully conserved across all marburgviruses, with the exception of RAVV GP for which 5 out of 15 residue positions differed. Since CM10, CM11, and CM12 all bind RAVV GP their binding either depends on conserved positions within this region or is agnostic to any sequence differences.

The third antigenic competition group we identified mapped to an epitope on GP1 that spanned residues 61-75, a partially conserved region across filoviruses. This region was previously identified as the target of a pan-filovirus reactive murine antibody m21D10 [39]. Four antibodies in our panel fell within this antigenic group, CM13, CM14, CM16, and CM16. While mAb CM13 recognized a peptide spanning this region as determined by ELISA, mAbs CM14, CM15, and CM16 did bind this peptide nor did they recognize denatured GP1. Nonetheless, binding competition assays revealed they could compete away roughly 40-50% of CM13 binding to RAVV GPΔMuc, suggesting they nonetheless target an overlapping epitope on GP1 but one that is likely conformational or complex.

Since animal NHP1 received a multivalent combination of filovirus immunogens, we also assessed the full antibody panel for pan-filovirus recognition of ebolavirus GPs. We found that two antibodies – CM16 and CM21 – both recognized multiple ebolavirus GPs including SUDV, EBOV, and BDBV. Indeed, their recognition of ebolavirus GPs appeared to exceed their capacity to recognize RAVV GPΔMuc. CM16 fell within the CM13 antigenic binding group, which as noted above was mapped to a conserved pan-filovirus epitope on GP1 overlapping that of murine antibody m21D10 [39]. Although CM21 did not fall within any of the characterized antigenic binding groups and could not be precisely mapped, we note that it did exhibit partial inhibition of CM13 binding to RAVV GPΔMuc in the competition assays, suggesting potential overlap of their epitopes nonetheless (**Fig. 6A,B**). The cross-filovirus recognition of GP detected for CM16 and CM21 thus confirms successful induction of antibody breadth.

Taken together, the present study yielded a novel panel of antibodies against marburgvirus GP isolated from a nonhuman primate immunized with an escalating species multivalent prime-boost regimen of filovirus antigens. Use of a heterologous probe based on RAVV GP ensured isolation of mAbs with cross-marburgvirus reactivity, including to our knowledge the first reported isolation of RBR-directed neutralizing antibodies from animal immunizations. While antibody neutralization capacity is a strong correlate of protective efficacy, further studies will be necessary to assess protective efficacy of the antibodies in animal challenge models. In addition, since 30% of mAbs in the panel could not be unambiguously mapped, further efforts will determine if they target novel conserved epitopes on the glycoprotein.

## Materials and Methods

### Antigen specific memory B cell sorting

Macaque monoclonal antibodies were isolated by single B-cell cloning as previously described [35, 43, 44]. In brief, macaque PBMCs were thawed and resuspend in staining media made up of RPMI 1640 supplemented with 10% fetal calf serum (FCS) at 37°. Cells were then washed in 10 mL staining media containing DNase I (Roche, Basel, Switzerland) and then resuspended in 100 μL of staining media containing 4 μg/mL of biotinylated RAVV GPΔMuc conjugated to streptavidin PE and 4 μg/mL of biotinylated RAVV GPΔMuc conjugated to streptavidin APC and incubated for 20 minutes [45]. This was followed by addition of a cocktail of CD3 APC-Cy7, CD8 APC-Cy7, Aqua Dead, CD14 Qdot 605 (BV605), IgM PE-Cy5, CD27 PE-Cy7, IgG FITC and CD20 PE-Alexa Fluor 700. Cells were gated for RAVV GPΔMuc double positive B-Cells with a phenotype CD27+, IgG+, CD20+, Aqua Dead-, CD3-, CD8-, IgM- and sorted at single cell precision on a BD FACSAria II [35]. Individual cells were sorted directly into lysis buffer and then subjected to reverse transcription-polymerase chain reaction (RT-PCR) using Superscript IV, as per manufacturer’s guidelines (Thermo Fisher Scientific, Waltham, MA). Nested PCR using HotStarTaq (Qiagen, Hildent, Germany) was then used amplify individual heavy and lambda/kappa light chains from the RT-PCR product. Heavy and light chain pairs were identified by agarose gel electrophoresis and sequenced by Sanger sequencing (Eurofins Genomics, Louisville, KY). Certain equipment, instruments, software, or materials, commercial or non-commercial, are identified in this paper in order to specify the experimental procedure adequately. Such identification is not intended to imply recommendation or endorsement of any product or service by NIST, nor is it intended to imply that the materials or equipment identified are necessarily the best available for the purpose.

### Expression of antibodies and GP proteins

Antibody variable heavy and light chain regions were synthesized by gene synthesis with appended N-terminal signal sequences (Genscript Biotech, Piscataway, NJ) and subcloned into IgG1 or lambda or kappa light chain based PCDNA3.1 mammalian expression plasmids. Plasmids were co-transfected into HEK-293F cells (ATCC, Manassas, VA) in FreeStyle Media using 293Fectin for transient protein expression (Thermo Fisher Scientific, Waltham, MA). Secreted IgGs were purified from cell supernatants with Protein A resin (Roche, Basel, Switzerland). IgGs were eluted at low pH using Protein A elution buffer (Thermo Fisher Scientific, Waltham, MA) and neutralized with Tris base pH 9.0. The IgGs were further purified by size exclusion chromatography (SEC) using an S200 column (Cytiva Lifesciences, Marlborough, MA) in PBS pH 7.4.

RAVV GPΔMuc fused with Strep II and Avi tags was expressed in HEK293S GNTI^−/−^ cells (ATCC, Manassas, VA) using 293fectin transfection reagent and FreeStyle Media (Thermo Fisher Scientific, Waltham, MA). Supernatants were purified using Streptactin XT Resin purification (IBA Lifesciences, Göttingen, Germany). The GP protein was further purified by SEC Superdex 200 HiLoad 16/600 column in 150 mM NaCl, 2.5 mM Tris-Cl pH 7.5 and 0.02% NaN3. GP was biotinylated by the Avi tag (Avidity, Aurora, Colorado) and exchanged into PBS 7.4.

### Pseudoparticle neutralization

Murine Leukemia Virus particles pseudotyped with GPs of EBOV, SUDV, BDBV, RESTV and MARV were prepared as previously described [46]. In brief, pseudoparticles were prepared by co-transfection of the human embryonic kidney 293T cells (HEK 293T) with the MLV Gag-Pol packaging vector (kindly provided by Dr. Jonathan K. Ball, University of Nottingham), a Luciferase reporter plasmid (kindly provided by Dr. Jonathan K. Ball), and plasmids of full-length GPs of five filoviruses corresponding to the following Genbank accession numbers: EBOV GP, AAN37507.1; SUDV GP, ALL26375.1; BDBV GP, AGL73460.1; RESTV GP, AAC24346.1; MARV GP, YP_001531156.1. Lipofectamine 3000 (Thermo Fisher Scientific, Waltham, MA) was used for these transfections, as per manufacturer’s protocols. A no-envelope control (empty plasmid) was the negative control in these experiments. At 6-hours post-transfection, the medium was replaced by fresh DMEM supplemented with 10% FBS. At 48-hours and 72-hours post-transfection, supernatants containing pseudoparticles were harvested, passed through 0.45 μm-pore-size filters, and then used to infect target cells. Luciferase activity was detected using BrightGlo (Promega, Madison, WI). Measured relative light units (RLU) were used to determine appropriate dilutions to use for the neutralization assays.

For the pseudoparticle neutralization assays, Vero E6 cells (ATCC, Manassas, VA) were pre-seeded into 96-wells plates at a density of 1×10^4^ per well. The following day, pseudoparticles were incubated with defined concentrations of mAbs for 1 hour at 37°C, and then added to each well of cells. Cells were then incubated in a CO_2_ incubator at 37°C for 5 hours. Mixtures on the cells were then replaced with fresh medium and cells were incubated for an additional 72 hours. Relative luciferase units (RLU) present in cell lysates was then assessed with BrightGlo using a FLUOstar Omega plate reader (BMG Labtech, Cary, NC) with the MARS software. All assays were performed in duplicate wells and tested plates contained both the positive and negative controls at the same dilutions. 50% inhibitory concentration titers (IC50s) were based on the concentrations of mAb that caused a 50% reduction in relative light units (RLU) compared with pseudoparticles in the control wells. Dose-response curves were fit with nonlinear regression plots in GraphPad Prism 7. Experiments using pseudoparticles were all performed under biosafety level 2 conditions.

### Pseudoparticle neutralization inhibition

The inhibition of the animal sera-mediated neutralization of MARV infection was tested using a neutralization inhibition assay in which the macaque NHP1 sera (study day 49 and pre-immune; [47]) were preincubated for 30 min with purified RAVV GPΔMuc at concentrations of either 2.5 μg/ml or 25 μg/ml before the addition of the MARV pseudoparticles. After incubating for 1 hour at 37°C, the mixtures were then added to the 96-well plates of Vero E6 cells and further incubated for 5 hours before replacing with fresh medium. With another 72-hour incubation, the luciferase activity was measured using the same method as described in the neutralization assays above. The inhibition effect of recombinant GP on MARV pseudoparticle neutralization was reported as the change between the serum ID50 with or without the presence of the tested GP competitor. The neutralization inhibition efficiency was calculated based on the following: [(percentage of neutralization w/o GPs-percentage of neutralization with GPs)/(percentage of neutralization w/o GPs)]x100. PBS was used as the negative control in the run.

### ELISA Assays

Nunc MaxiSorp 96 well ELISA plates (Thermo Fisher Scientific, Waltham, MA) were coated at 4°C overnight with filovirus GPs. The plates were washed in wash buffer PBS 7.4 containing 0.05% Tween 20 and then blocked in PBS pH 7.4, 5% fetal bovine serum and 2% non-fat dry milk powder for one hour at room temperature. The plates were washed and a series of 5-fold serial dilutions of monoclonal antibody starting at 10 μg/mL added. Plates were incubated for one hour, washed, and then incubated with a 1:2500 dilution of horseradish peroxidase-conjugated goat anti-human secondary antibody (Jackson ImmunoResearch, West Grove, PA) in blocking buffer for one hour. The plates were developed with TMB ELISA substrate solution (Bio-Rad Laboratories, Hercules, CA) and stopped using 1 N sulfuric acid. Plates were read at an absorbance of 450 nm.

Competition ELISAs were undertaken by coating half area ELISA plates (Greiner Bio-One, Monroe, NC) with 1 µg/mL of the benchmark antibodies at 4°C overnight. The next day plates were washed in wash buffer containing PBS pH 7.4 supplemented with 0.05% Tween 20 and then blocked in PBS pH 7.4, 5% fetal bovine serum, and 2% non-fat dry milk powder for one hour at room temperature. During this time in a separate non-binding U well shape plate (Greiner Bio-One, Monroe, NC) competing antibody was diluted to a final concentration of 5 µg/mL in blocking buffer and added to 2 µg/mL GP that was biotinylated through a fused Avi-tag (Avidity, Aurora, Colorado) and incubated for one hour at room temperature. The GP antibody mixture was then added to ELISA plates coated with capture antibody and incubated for 1 hour at room temperature. Plates were washed in wash buffer and then incubated with a 1:10,000 dilution of goat anti-biotin antibody (Thermo Fisher Scientific, Waltham, MA) in PBS pH 7.4 supplemented with 0.05% tween-20. The ELISA was developed as described above.

### Pepscan analysis

Peptide competitions were undertaken by incubating RAVV GP ΔMuc at 200 ng per a well in Nunc MaxiSorp 96 well ELISA plates (Thermo Fisher Scientific, Waltham, MA) overnight at 4°C. The plates were blocked as above. In separate non-binding U-well shaped plates (Greiner Bio-One, Monroe, NC) 200 ng of each of the overlapping 15-mer peptides across either GP1 or GP2 was incubated individually with 0.4 µg/mL of IgG for 1 hour at room temperature. 100 µL of each IgG peptide mixture was added to the plate with GP and incubated for 1 hour. Plates were washed and a horseradish peroxidase-conjugated goat anti-human secondary antibody (Jackson ImmunoResearch, PA) was added at a 1:2500 dilution. Plates were washed and developed as above. Direct pepscan analysis was undertaken by binding 200 ng of each overlapping peptide to Nunc MaxiSorp 96 well ELISA plates (Thermo Fisher Scientific, Waltham, MA) overnight at 4°C. Plates were washed and blocked as above, and 0.4 µg/mL of IgG was then added to each well for 1 hour at room temperature. Plates were washed and secondary antibody added as described above. Finally, plates were washed and developed as described above.

### Western blots

Mini-PROTEAN TGX gels were run and transferred to a PVDF or nitrocellulose membrane using the Turbo Blot protocol for mini-PROTEAN TGX gels (Bio-Rad Laboratories, Hercules, CA). The membranes were blocked for 5 min with 5% skim milk powder. The PVDF/nitrocellulose membranes were then divided into strips of one lane, each containing GP1 and GP2. The strips were placed in separate primary stain solutions with each individual antibody at 1 µg/mL. After staining on a platform rocker for 1 hour, the strips were washed 3 times for five minutes with TBST. All strips were then stained separately with a 1:5000 dilution of goat anti-human IgG for 1 hour. After the secondary staining, the strips were again washed 3 times with TBST for five minutes. The strips were then placed together and developed by enhanced chemiluminescence (ECL) (Thermo Fisher Scientific, Waltham, MA) and imaged on a ChemiDoc imager (Bio-Rad Laboratories, Hercules, CA).

### Negative stain electron microscopy

Negative staining was performed following the optimized negative staining (OpNS) protocol as described [48]. Optimized negative staining: a high-throughput protocol for examining small and asymmetric protein structure by electron microscopy [48]. Briefly, complexes were diluted to 0.01 mg/mL and immediately applied to EM grids (Electron Microscopy Sciences #CF200-Cu). Grids were then incubated for one minute, blotted with filter paper, washed three times with water as described, and stained with fresh 1% uranyl formate solution for 30 s as described. Staining solution was then blotted with filter paper and grids were dried in a desiccator overnight prior to imaging.

Imaging was performed using a Talos Arctica (200 kV) system (Thermo Fisher Scientific) equipped with a Falcon 3EC detector. A nominal magnification of 73,000x was used, corresponding to a pixel size of 1.38 Å. Dose-fractionated movies were collected with a total dose of about 120 e/Å^2^, and motion correction was performed using RELION [41]. Particle picking was done using crYOLO followed by 2D classification in RELION [41, 49].

## Author Contributions

Conceptualization, T.R.F. and G.O.; Formal Analysis, B.M.J., R.W., T.E.C.; Investigation, B.M.J., R.W, T.E.C., M.C.M., A.C.L., K.G., K.C., and G.O.; Methodology, B.M.J., R.W., T.E.C., T.R.F., and G.O.; Project Administration, G.O.; Supervision, T.R.F. and G.O; Visualization, B.M.J., R.W, T.E.C., and G.O.; Writing – Original Draft Preparation, G.O.; Writing – Review & Editing, B.M.J., R.W, T.E.C., M.C.M, A.C.L., K.C., T.R.F., and G.O.; Funding Acquisition, T.R.F. and G.O.

